# Vector acquisition and co-inoculation of two plant viruses influences transmission, infection, and replication in new hosts

**DOI:** 10.1101/2022.08.27.505557

**Authors:** Autumn A. McLaughlin, Linda Hanley-Bowdoin, George G. Kennedy, Alana L. Jacobson

**Affiliations:** Department of Entomology and Plant Pathology, Auburn University, Auburn, AL 36849; Department of Plant and Microbial Biology, North Carolina State University, Raleigh, NC 27695; Department of Entomology and Plant Pathology, North Carolina State University, Raleigh, NC 27695

## Abstract

This study investigated the role of vector acquisition and transmission on the propagation of single and co-infections of tomato yellow leaf curl virus (TYLCV,) and tomato mottle virus (ToMoV) (Family: *Geminiviridae*, Genus: *Begomovirus*) by the whitefly vector *Bemisia tabaci* MEAM1 (Gennadius) in tomato. The aim of this research was to determine if how viruses are co-acquired and co-transmitted changes the probability of acquisition, transmission and new host infections. Whiteflies acquired virus by feeding on singly infected plants, co-infected plants, or by sequential feeding on singly infected plants. Viral titers were also quantified by qPCR in vector cohorts, in artificial diet, and plants after exposure to viruliferous vectors. Differences in transmission, infection status of plants, and titers of TYLCV and ToMoV were observed among treatments. All vector cohorts acquired both viruses, but co-acquisition/co-inoculation generally reduced transmission of both viruses as single and mixed infections. Co-inoculation of viruses by the vector also altered virus accumulation in plants regardless of whether one or both viruses were propagated in new hosts. These findings highlight the complex nature of vector-virus-plant interactions that influence the spread and replication of viruses as single and co-infections.

## Introduction

Concurrent infections of two or more plant viruses, referred to as co-infections or mixed infections, are common in nature and are known to influence disease severity, spread, and the emergence of novel variants^1^. Fitness of the co-infecting viruses may be altered by virus-virus and virus-plant interactions that change virus replication, accumulation, movement, and tissue tropism in the plant^2–8^. These can influence foliar symptom severity, growth, development, and chemical and physiological profiles of the infected plants^2,3,9^ that can alter vector behavior in ways that affect virus acquisition and transmission^3–6,10–22^. Most research to-date has been conducted on single pathogen systems, and knowledge of more complex co-infection dynamics is limited. A majority of studies on mixed infections examine within plant-host dynamics of two viruses and their outcomes on disease severity and viral fitness^2–8^. Comparatively few studies use insect vector-transmission or examine vector-related effects^10,11,22–41^. This study was conducted to examine how the manner of acquisition and inoculation of two viruses by their common vector affects propagation of co-infections in their plant hosts.

Co-infections have both spatial and temporal dimensions based on when and where plant inoculation occurs^8,42–46^. Plant hosts are spatial environments that viruses must effectively exploit to replicate and localize in tissue so that virions are acquired by their vectors^20,21,47^. Co-infections of plant viruses can be propagated by vectors that acquired virions from co-infected plants, sequentially from singly infected plants, or by multiple vector individuals that acquired different viruses. Vector acquisition of two viruses does not ensure both will be transmitted, and the presence of two viruses can alter transmission rates of one or both viruses^4,10,11,22–32,34–41,48–51^. Once acquired, two viruses may compete for binding sites required for retention in and transmission from their vectors^6,29,33,37,49,52^. Order of acquisition sometimes has an antagonistic effect on virus transmission^4,10,34,36,38^ and does not always benefit the virus acquired first^22,30,36^. Vector-plant interactions that occur during probing and feeding events can also induce host plant defenses that mediate infection dynamics.

This study was undertaken to better understand how vector acquisition and transmission of two viruses influences virus persistence. Tomato yellow leaf curl virus (TYLCY) and tomato mottle virus (ToMoV) (Family: *Geminiviridae*, Genus: *Begomovirus*) are members of the largest plant virus genus for which the importance of co-infections has been documented as a key feature of the emergence, severity and evolution of economically important species^53–55^. Both are transmitted in a persistent manner by whiteflies in the *Bemisia tabaci* (Gennadius) cryptic species complex and occur in single and co-infections in tomato production systems in the southeastern U.S.A^56,57^. Begomovirus genomes are circular single-stranded DNA (ssDNA) molecules of approximately 2800 nucleotides each that are encapsulated separately in a 22 × 38 nm geminate particles^58–60^. ToMoV has a bipartite genome comprised of two ssDNA components, DNA-A and DNA-B, whereas TYLCV is monopartite with all of its genes encoded on one ssDNA molecule. Both components of bipartite begomoviruses are required to initiate an infection because DNA-A encodes proteins necessary for viral replication, encapsidation, transcription and plant defense neutralization, while DNA-B encodes the viral movement proteins^61,62^.

To test the hypothesis that virus-virus interactions involving the vector and host plants change viral fitness to be acquired from, transmitted and propagated within new hosts, a transmission experiment was designed to investigate the effects of virus acquisition scenarios on 1) probability of virus acquisition by the vector, 2) virus titers in the vector after acquisition, 3) probability of virus transmission, 4) titers of virus transmitted, 5) probability of systemic infection in inoculated host plants, and 6) virus titers of newly infected host plants (Table 1). Virus detection in whitefly, artificial diet and plant samples provided measures of the probability of vector acquisition, vector transmission, and new host infection in each treatment. Quantification of virus titers provided information on virus accumulation in vectors, artificial media after transmission, and replication in systemically infected hosts. This study provides new knowledge about the importance of virus-virus-plant, virus-virus-vector, and vector-plant interactions on the propagation of two viruses as single and co-infections.

**Table 1.** Treatments in this experiment were designed to compare acquisition and transmission of single and co-infections TYLCV and ToMoV under different acquisition and inoculation scenario treatments.

| **Treatment** | **Virus Acquisition Scenarios** | **Virus Inoculation Scenarios** |
| --- | --- | --- |
| <u>Singly Infected sources:</u><br>1. TYLCV<br>2. ToMoV | Acquisition of a single virus; 48 h acquisition access period (AAP) on singly infected plants. | Inoculation of a single virus. |
| <u>Co-infected source:</u><br>3. Mixed | Acquisition of two viruses from one plant co-infected with TYLCV and ToMoV; 48 h AAP | Co-inoculation may occur via an individual. |
| <u>Sequential Acquisitions:</u><br>4. TYLCV»ToMoV<br>5. ToMoV»TYLCV | Acquisition of two viruses occurs sequentially from two singly infected plants; 48 h AAP total, 24 h on each infected plant. |  |
| <u>Mixed Vector Cohorts:</u><br>6. TYLCV+ToMoV | Acquisition of each virus by different vector cohorts combined in equal numbers for a 48 h AAP on singly infected plants. | Co-inoculation requires at least two individuals; one from the TYLCV cohort and one from the ToMoV cohort. |
| 7. Healthy Control | Whiteflies feed on a healthy, uninfected source plant. | Check for contamination of whiteflies and cross-contamination in incubators. |

## Results

The experimental treatments included two single infection controls and four scenarios involving acquisition of two viruses by their vectors (Table 1). No standard terminology exists for describing the co-acquisition and co-transmission processes, or vector-related spatial and temporal dimensions of the processes. In our experiments, “co-acquisition” refers to vector acquisition of two viruses from one co-infected plant, whereas “sequential acquisition” refers to vector acquisition of two viruses from two sequential acquisition access periods (AAP) on singly infected hosts. “Co-inoculation” refers to virus inoculation occurring during the same inoculation access period (IAP) after the vector co-acquires both viruses. “Mixed cohort scenario” refers to a group of viruliferous vectors in which individuals acquired different viruses. “Transmission” refers to virus detection in an artificial diet after vector feeding, and represents a measure of transmission that is not confounded by replication in host plants or host defenses. “Infection” refers to systemic virus infections in vector-inoculated plants and reflects the outcome of vectorvirus-plant interactions that occurred during and after vector co-inoculation. In these experiments, both “co-infections” of ToMoV and TYLCV and single infections of each virus were detected after vector co-inoculation. The term “vector-co-inoculated, singly-infected plants” is used to distinguish single infections propagated by vectors shown to co-acquire both viruses in the Mixed, TYLCV»ToMoV, ToMoV»TYLCV, and TYLCV+ToMoV treatments from single infections used in the TYLCV and ToMoV control treatments. Because ToMoV is bipartite and both DNA-A and DNA-B must be transmitted to the plant for infection to occur, the terms acquisition and transmission when applied to ToMoV indicate both components were detected.

### Virus Titers of Infected Plants Used for AAPs

To standardize virus populations in these experiments *Rhizobacterium tumenfacious* infectious clones of TYLCV and ToMoV^63^ were used to initiate infections in *Solanum lycopersicum* L., variety ‘Florida Lanai^64,65^ (rhizobacteriuminoculated plants); these plants were used for virus acquisition during the AAPs. There were significant differences in the titers of TYLCV and ToMoV DNA-components in rhizobacterium-inoculated plants, but the titers of each viral component reached similar levels in singly and coinfected plants (*P*<0.0001, Supplemental Figure 1). TYLCV titers were the highest, but not statistically higher than ToMoV-B co-infected plants, whereas ToMoV-A was lower than both TYLCV and ToMoV-B.

### Quantifying Virus Acquisition by Whiteflies

Whitefly cohorts efficiently acquired viruses from source plants regardless of the acquisition scenario. Five separate analyses compared the proportion of cohorts that acquired one virus (TYLCV only, ToMoV only), both viruses (TYLCV & ToMoV), or the total of single and co-acquisitions (Total TYLCV, Total ToMoV) among treatment scenarios (Table 2). There were no significant differences in acquisition among treatments. Both viruses were detected in all but one sample from the co-acquisition scenario treatments, but there was not enough DNA to re-run the one sample to retest.

**Table 2.**
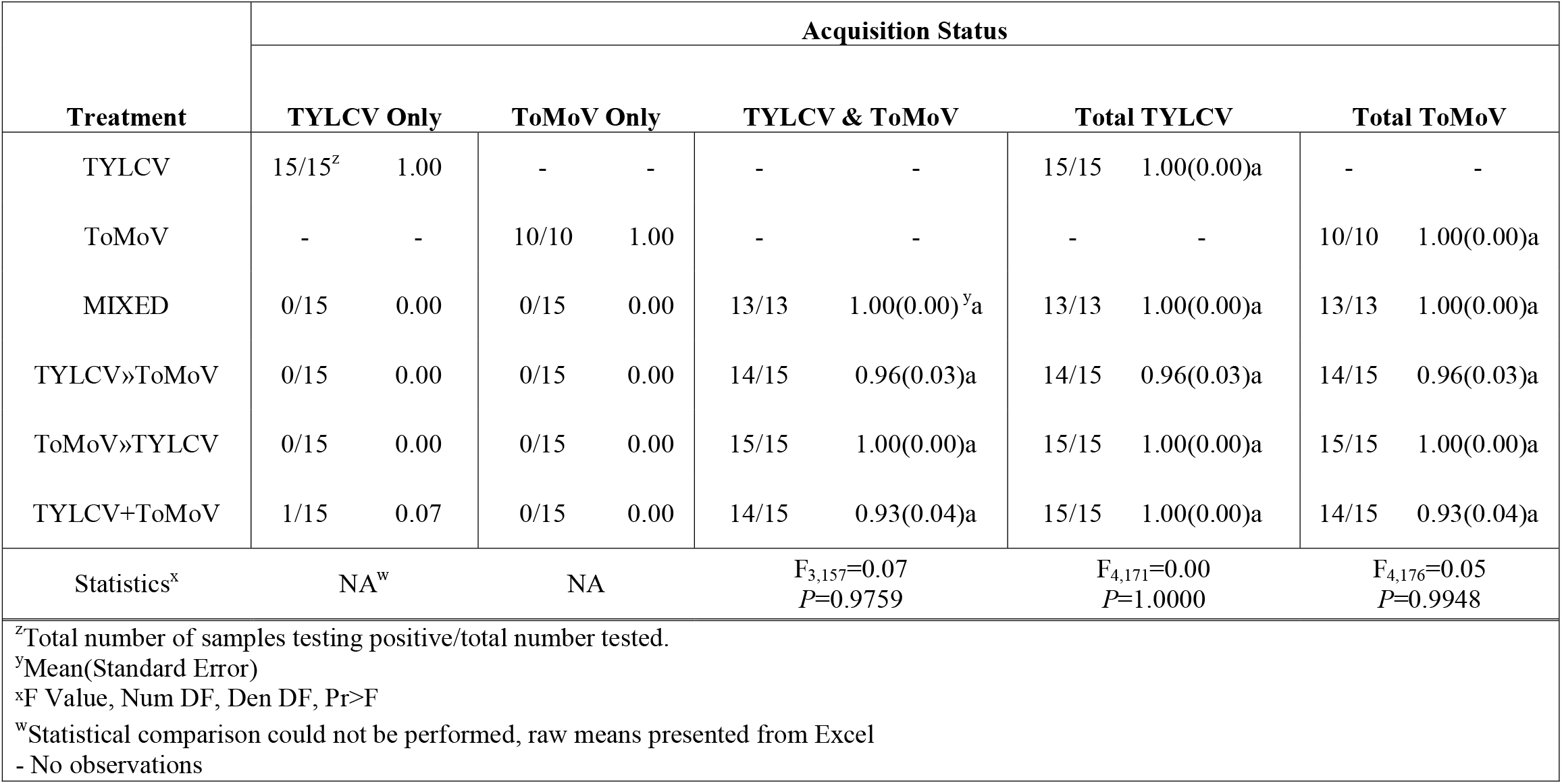
The average proportion of whitefly cohorts that acquired TYLCV or/and ToMoV. LS means comparisons among treatments were conducted using Tukey’s method at *P*=0.05 level to compare acquisition of viruses singly (TYLCV only, ToMoV only columns), together (TYLCV & ToMoV column), and sum of virus infection (Total TYLCV, Total ToMoV columns).

Average virus titer in the whitefly cohorts was measured using qPCR. The main effect of acquisition scenario treatment was tested in three separate analyses on TYLCV, ToMoV-A and ToMoV-B that included single and co-infections (Figure 1A). There were no significant differences among treatments for TYLCV (F_4,65_=1.76, *P=*0.1476), ToMoV-A (F_4,60_=1.64, *P=*0.1761), or ToMoV-B (F_4,60_=1.13, *P=*0.3492). An additional analysis on virus titers included only whitefly cohorts that had acquired both viruses from co-acquisition scenario treatments (no single infection treatments or samples that tested positive for only one virus) to compare the main effects of treatment, virus/viral component, and their interaction. There were no significant differences among treatments (F_3,153_=2.20, *P=*0.0904; Supplemental Figure 2A) or the treatmentby-virus/viral component interaction (F_6,153_=0.65, *P*=0.6917; Figure 2A), but the titer of TYLCV acquired by whiteflies, averaged across treatments, was significantly higher than the titers of ToMoV-A and ToMoV-B (F_2,153_=14.67, *P<*0.0001 ; Supplemental Figure 2B).

**Figure 1.**
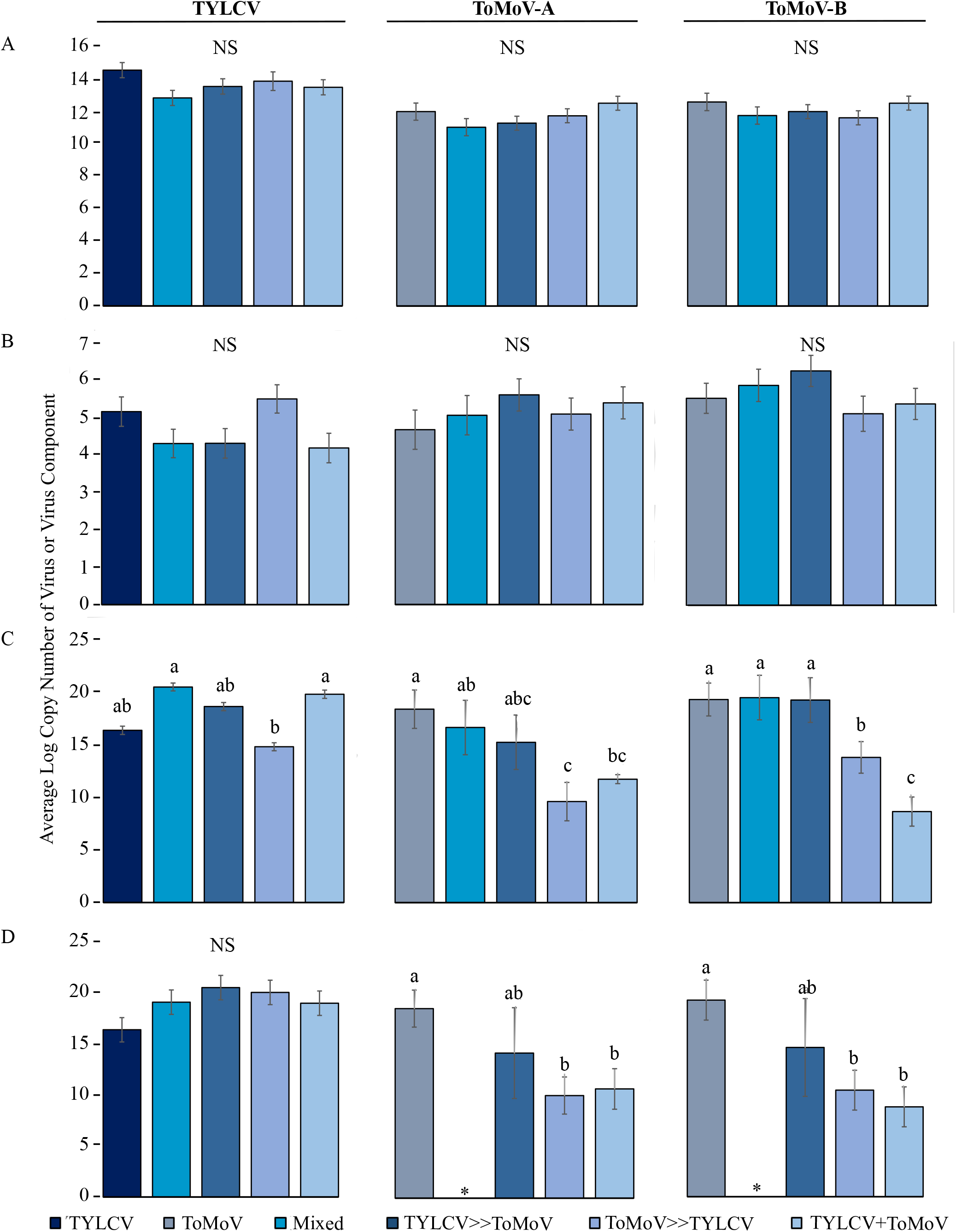
The average log_10_ copy number of TYLCV, ToMoV-A, and ToMoV-B were quantified and analyzed separately in (A) vector cohorts; (B) artificial diet viruliferous vector cohorts fed on (C) singly infected control plants (TYLCV or ToMoV treatments) and plants coinfected after vector inoculation in co-infection scenario treatments (Mixed, TYLCV»ToMoV, ToMoV»TYLCV, TYLCV+ToMoV), and (D) singly infected control plants and vector-(co-) inoculated singly-infected plants (VCSIPs). *No single infections of ToMoV occurred as VCSIPs in the Mixed treatment. Copy numbers were compared for the main effects of virus acquisition scenario treatment, virus or viral component, and their interaction term; averages from interaction term main effects are shown. LS means comparisons were conducted using Tukey’s method at *P*=0.05. NS represents that the interaction term in the model was not significant.

**Figure 2.**
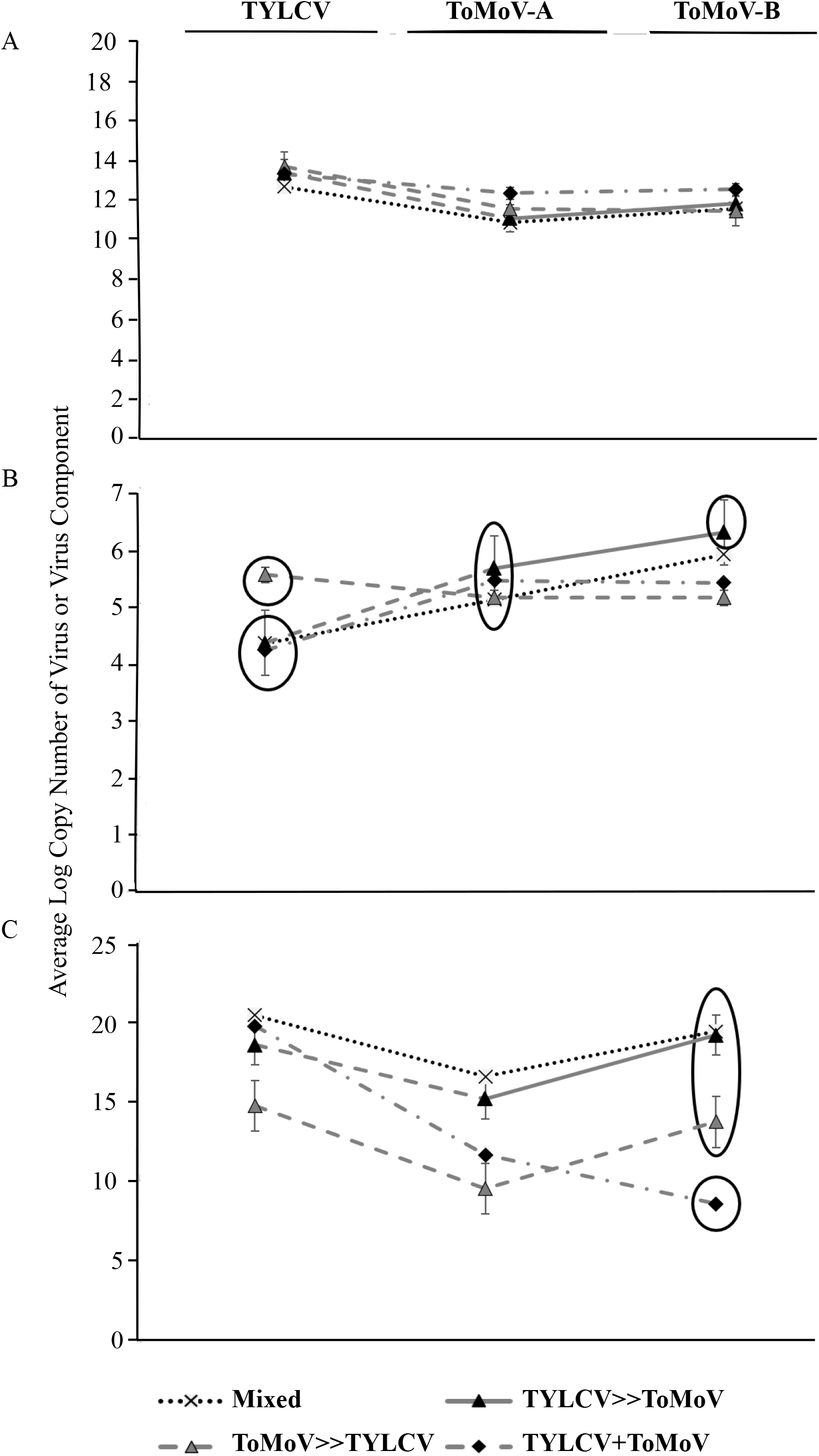
The average log_10_ copy number of TYLCV, ToMoV-A, and ToMoV-B in co-acquisition scenario treatments. Data were analyzed separately for samples in which both TYLCV and ToMoV were detected in (A) vector cohorts, (B) artificial diet viruliferous vector cohorts fed upon, and (C) co-infected plants inoculated with vector cohorts. Copy numbers were compared for the main effects of virus acquisition scenario treatment, virus or viral component, and their interaction term; averages from interaction term main effects are shown. LS means comparisons were conducted using Tukey’s method at *P*=0.05. Circled averages are not significantly different from each other by virus/viral component; no circles indicate no differences.

### Quantifying Virus Transmission to Artificial Diet

Virus transmission efficiency was examined by analyzing the proportion of sucrose sachets (artificial diet packets) containing only one virus (TYLCV only or ToMoV only), both viruses (TYLCV & ToMoV), and the sum of single and co-transmissions (Total TYLCV, Total ToMoV) in separate analyses (Table 3). Although virtually all whitefly cohorts acquired both viruses in co-acquisition scenario treatments, very few transmitted both viruses to sucrose sachets. The transmission efficiency of TYLCV singly differed significantly among treatments. TYLCV transmission was lower when the whitefly cohorts co-acquired both viruses (Mixed), acquired TYLCV prior to ToMoV (TYLCV»ToMoV), or in the mixed cohort scenario treatment (TYLCV+ToMoV) compared to the single infection scenario. But when TYLCV was acquired by the whitefly cohort following acquisition of ToMoV (ToMoV»TYLCV), transmission efficiency was comparable to the single infection scenario. In contrast, transmission of ToMoV (ToMoV-A and ToMoV-B) in the Mixed treatment was not significantly different from the single infection scenario. Transmission of ToMoV was significantly reduced when both viruses were acquired sequentially by the whitefly cohorts (TYLCV»ToMoV, ToMoV»TYLCV), and in the mixed cohort scenario (TYLCV+ToMoV). Transmission of both viruses by whitefly cohorts was low (0 – 13%) and not affected by acquisition scenario.

**Table 3.** The average proportion of sucrose sachets inoculated with TYLCV or/and (ToMoV) by whitefly cohorts. LS means comparisons among treatments were conducted using Tukey’s method at *P*=0.05 level to compare the inoculation of viruses singly (TYLCV only, ToMoV only columns), together (TYLCV & ToMoV column), and sum of virus infection (Total TYLCV, Total ToMoV columns).

|  | Transmission Status |  |  |  |  |  |  |  |  |  |
| --- | --- | --- | --- | --- | --- | --- | --- | --- | --- | --- |
| Treatment<br>Treatment | Infection Status |  |  |  |  |  |  |  |  |  |
|  | TYLCV Only |  | ToMoV Only |  | TYLCV & ToMoV |  | Total TYLCV |  | Total ToMoV |  |
| TYLCV | 9/15<sup>z</sup> | 0.60(0.12)<sup>y</sup>a | - | - | - | - | 9/15 | 0.60(0.13)ab | - | - |
| ToMoV | - | - | 7/15 | 0.47(0.13)a | - | - | - | - | 7/15 | 0.47(0.09)a |
| MIXED | 4/15 | 0.29(0.07)b | 5/14 | 0.36(0.09)ab | 0/15 | 0.00(0.00)a | 4/15 | 0.29(0.07)c | 5/15 | 0.36(0.07)ab |
| TYLCV»ToMoV | 4/15 | 0.27(0.07)bc | 0/15 | 0.00(0.00)c | 1/15 | 0.06(0.03)a | 5/15 | 0.33(0.07)bc | 1/15 | 0.07(0.04)c |
| ToMoV»TYLCV | 9/15 | 0.60(0.07)a | 1/15 | 0.06(0.05)c | 2/15 | 0.13(0.05)a | 11/15 | 0.73(0.07)a | 3/15 | 0.20(0.06)bc |
| TYLCV+ToMoV | 2/15 | 0.11(0.05)c | 2/15 | 0.13(0.06)bc | 2/15 | 0.13(0.05)a | 4/15 | 0.27(0.07)c | 4/15 | 0.27(0.07)ab |
| Statistics<sup>x</sup> | F<sub>4,187</sub>=6.22,<br>P<0.0001 |  | F<sub>4,128</sub>=2.89,<br>P=0.0251 |  | F<sub>3,173</sub>=0.43, P=0.7294 |  | F<sub>4,187</sub>=6.53, P<0.0001 |  | F<sub>4,202</sub>=3.90, P=0.0045 |  |
| <sup>z</sup>Total number of samples testing positive/total number tested.<br><sup>y</sup>Mean(Standard Error)<br><sup>x</sup>F Value, Num DF, Den DF, Pr>F<br>-No observations |  |  |  |  |  |  |  |  |  |  |

The average quantity of virus transmitted by the whitefly cohorts to artificial diet was measured with qPCR. The main effect of acquisition scenario treatment was analyzed separately for TYLCV, ToMoV-A and ToMoV-B, and included single and co-infections (Figure 1B). There were no significant differences among treatments for TYLCV (F_4,61_=2.38, *P=*0.0612), ToMoVA (F_4,62_=1.32, *P=*0.2721), and ToMoV-B (F_4,61_=1.06, *P=*0.3862). An analysis of virus titers that included only cohorts that transmitted both viruses from co-acquisition scenario treatments revealed the main effect of acquisition scenario treatment was not significant (F_3,149_=0.73, *P=*0.5368; Supplemental Figure 3A). However, there were significant differences in virus/viral component titers (F_2,149_=8.57, *P<*0.0003; Supplemental Figure 3B); ToMoV-A and ToMoV-B titers were significantly higher than TYLCV. The interaction term was also significant (F_6,149_=2.21, *P*=0.0451; Figure 2B), indicating that differences in virus titers among treatments depended on the virus component.

### Transmission to Plants

Separate analyses compared the proportion of vector-inoculated plants that became systemically infected with one virus (TYLCV only, ToMoV only), both viruses (TYLCV & ToMoV), and the total of single and co-infections (Total TYLCV, Total ToMoV) (Table 4). There were significant differences in the number of singly infected plants, but not in the number of co-infected plants among treatments. The proportion of plants infected with TYLCV or ToMoV in the co-acquisition scenario treatments was generally reduced compared to the single infection treatments.

**Table 4.** The average proportion of vector inoculated plants systemically infected with TYLCV or/and ToMoV. LS means comparisons among treatments were conducted using Tukey’s method at *P*=0.05 level to compare infection of viruses singly (TYLCV only, ToMoV only columns), together (TYLCV & ToMoV column), and sum of virus infection (Total TYLCV, Total ToMoV columns).

|  | TYLCV Only |  | ToMoV Only |  | TYLCV & ToMoV |  | Total TYLCV |  | Total ToMoV |  |
| --- | --- | --- | --- | --- | --- | --- | --- | --- | --- | --- |
| TYLCV | 37/51<sup>z</sup> | 0.73(0.06)<sup>y</sup>a | - | - | - | - | 37/51 | 0.73(0.06)a | - | - |
| ToMoV | - | - | 17/51 | 0.30(0.06)a | - | - | - | - | 17/51 | 0.30(0.06)a |
| MIXED | 18/43 | 0.42(0.08)bc | 0/43 | 0.00(0.00)c | 3/43 | 0.07(0.03)a | 21/43 | 0.48(0.08)b | 3/43 | 0.07(0.04)b |
| TYLCV»ToMoV | 23/49 | 0.45(0.07)b | 1/49 | 0.02(0.02)c | 3/49 | 0.06(0.03)a | 26/49 | 0.53(0.07)b | 4/49 | 0.10(0.03)b |
| ToMoV»TYLCV | 10/46 | 0.22(0.06)c | 11/46 | 0.24(0.06)ab | 6/46 | 0.13(0.06)a | 16/46 | 0.36(0.07)b | 17/46 | 0.38(0.07)a |
| TYLCV+ToMoV | 12/41 | 0.29(0.07)bc | 5/41 | 0.12(0.05)bc | 8/41 | 0.20(0.07)a | 20/41 | 0.48(0.08)b | 13/41 | 0.32(0.07)a |
| Statistics<sup>x</sup> | F<sub>4,214</sub>=6.61,<br>P<0.0001 |  | F<sub>4,230</sub>=2.75,<br>P=0.0289 |  | F<sub>3,174</sub>=1.57,<br>P=0.1978 |  | F<sub>4,214</sub>=3.28,<br>P=0.0122 |  | F<sub>4,225</sub>=4.16,<br>P=0.0029 |  |
| <sup>z</sup>Total number of samples testing positive/total number tested. |  |  |  |  |  |  |  |  |  |  |
| <sup>y</sup>Mean(Standard Error) |  |  |  |  |  |  |  |  |  |  |
| <sup>x</sup>F Value, Num DF, Den DF, Pr>F |  |  |  |  |  |  |  |  |  |  |
| -No observations |  |  |  |  |  |  |  |  |  |  |

### Virus Accumulation in Co-infected Plants

The main effect of acquisition scenario treatment on virus and viral component titers was analyzed separately for TYLCV, ToMoV-A and ToMoV-B, and included the single infection control treatments. There were significant differences among treatments in the accumulation of TYLCV (F_4,20_=2.90, *P=*0.0481), ToMoV-A (F_4,18_=3.53, *P=*0.0271), and ToMoV-B (F_4,20_=9.58, *P=*0.0002) (Figure 1C). TYLCV titers in plants did not differ significantly between the single infection treatment and any of the coinfection treatments. ToMoV-A and ToMoV-B titers were significantly lower when cohorts acquired ToMoV first (ToMoV»TYLCV) and in the mixed cohort scenario (TYLCV+ToMoV).

In the analysis that included only co-acquisition scenario treatments, there were significant differences in the main effects of treatment (F_3, 43_=7.10, *P=*0.0006; Supplemental Figure 4A), virus or viral component (F_2, 43_=6.74, *P*=0.0029; Supplemental Figure 4B), and their interaction (F_6, 43_=2.71, *P*=0.0254; Figure 2C). The average accumulation of TYLCV across all treatments was significantly higher than both ToMoV-A and ToMoV-B (Supplemental Figure 4B). The significant interaction term reflects similar titers of TYLCV or ToMoV-A among treatments, but a significantly lower titer of ToMoV-B when ToMoV was acquired by different whiteflies than TYLCV (TYLCV+ToMoV).

### Virus Accumulation in Vector-Co-inoculated, Singly-Infected Plants (VCSIPs)

Differences in virus or viral component titers among treatments were analyzed separately for TYLCV, ToMoV-A and ToMoV-B in VCSIPs (Figure 1D). There were no differences in TYLCV accumulation among treatments (F_4, 25_=1.79, *P=*0.1622), but there were differences in ToMoV-A (F_3, 14_=4.46, *P=*0.0213) and ToMoV-B (F_3, 15_=5.56, *P=*0.0091). Accumulation of ToMoV components were lower when ToMoV was acquired first (ToMoV»TYLCV) or separately from TYLCV (TYLCV+ToMoV) compared to the ToMoV treatment. Accumulation was not statistically reduced in the TYLCV»ToMoV treatment, but the sample size was low (only two ToMoV-VCSIPs were observed in this treatment). No ToMoV-VCSIPs were observed in the Mixed treatment.

## Discussion

Our results provide new insights into the effects of virus co-acquisition and coinoculation on (co-)infection of new host plants and contribute to the expanding body of research on how viruses interact directly and indirectly with other viruses, their vectors, and hosts to affect acquisition, spread and accumulation. In this study, the order of virus acquisition did not change the probability of acquisition by vector cohorts (Table 2) but did influence transmission efficiency of the viruses (Table 3) and the probability of systemic host plant infection (Table 4). Co-infections were propagated in all co-acquisition scenario treatments, but their occurrence was very low due to antagonistic virus-virus interactions in the vector and/or host plants. These interactions also changed the accumulation of viruses that persisted as single and co-infections (Figures 1C-D, 2C).

### Vector acquisition and transmission of two viruses

Although less commonly described in the context of fitness outcomes, the probability of vector acquisition and transmission to new hosts are components of virus fitness mediated by the vector. The amounts of virus ingested by a vector may vary for different viruses^66^, can be influenced by where viruses localize in plant tissues^5,10,13–15^, and the titers of virus that accumulate in those tissues^10,67^. Titers of TYLCV were higher than ToMoV in single and co-infected rhizobacterium-inoculated plants used in the AAPs (Supplemental Figure 1), but there is not strong evidence that higher titers of TYLCV impaired vector acquisition (Table 2) and/or transmission (Table 3) of ToMoV. Both viruses were detected in almost 100% of vector cohorts, and although when averaged across all treatments the titer of TYLCV in whitefly cohorts was higher than ToMoV, their respective titers within each acquisition scenario treatment were not significantly different. (Figure 1A). Replication of TYLCV in *B. tabaci* could contribute to the higher average titers of TYLCV in vector cohorts^68–70^, but replication is not reported to occur for all TYLCV-isolate x *B. tabaci* population combinations^71–73^, and measuring replication of TYLCV and ToMoV in vectors was outside the scope of this study. Importantly, although whiteflies in the sequential acquisition scenarios had a 24 h AAP for each virus while those in the other acquisition scenarios had a 48 AAP, this difference did not affect virus titers in the whiteflies and does not explain effect of the sequential treatment on transmission rates of either virus.

Factors associated with virus co-circulation through vectors influenced transmission outcomes. The relative titers of TYLCV and ToMoV detected in artificial diet did not reflect titers in the corresponding transmitting whitefly cohorts or the plants from which they acquired virus. Averaged across treatments, titers of ToMoV-A and ToMoV-B in artificial diet were higher than TYLCV (Supplemental Figure 3B), and the average ToMoV component titers by treatment were significantly higher in all treatments except TYLCV+ToMoV (Figure 1B). This suggests that although this isolate of TYLCV had an overall higher fitness in plants, the ToMoV isolate was generally better able to circulate through the vector and be transmitted. However, despite higher average titers of ToMoV components, the transmission of both viruses to artificial diet was reduced in most co-acquisition scenario treatments (Table 2).

The set of optimal within-vector interactions that maximize viral fitness will depend largely on the mode of transmission of the virus and may be altered by co-infections^19,74–76^. Begomoviruses are subject to physiological, immunological, and endosymbiont-related effects within their vector that may influence retention, circulation and transmission^77–81^. Previous studies on vector transmission of two viruses focused mainly on non-persistently and semipersistently transmitted viruses, and their findings regarding the effect of acquisition scenario on transmission are inconsistent. Our study examined persistently transmitted viruses that move across the vector gut, circulate in the hemolymph, and localize in salivary ducts from which transmission occurs. The vector cohort size and acquisition and inoculation access times used were chosen to maximize transmission rates based on previously published studies^82–85^.

Within our experimental design, the virus acquired first in co-acquisition scenario treatments (virus_1_), should be transmitted first because it had 24 h to circulate in the vector before acquisition of virus acquired second (virus_2_). All else being equal, this is expected to result in the predominance of virus_1_ at initial inoculation sites in plants during the first 24 hours of the IAP. If the viruses compete within the vectors for binding sites required for retention and circulation, then sequential acquisition scenarios should also favor virus_1_. This effect would be greater if initial vector interactions with virus_1_ produce an antagonistic effect on circulation of virus_2_. It is also possible, however, that vector interactions with virus_2_ may result in antagonism on virus_1_, or that vector-virus_1_-virus_2_ interactions may produce antagonistic effects for both viruses. In the Mixed treatment the order of acquisition from the co-infected plant presumably varied among individual whiteflies within a cohort. The acquisition of both viruses reduced the transmission of TYLCV, but not ToMoV to artificial diet (Table 3). In contrast, sequential acquisition, regardless of the order in which the viruses were acquired, reduced transmission of ToMoV but not TYLCV to artificial diet (Table 3). When individual whiteflies in the transmitting cohort acquired only TYLCV or ToMoV (TYLCV+ToMoV treatment) total transmission to artificial diet were reduced for TYLCV but not ToMoV. Because there is no consistent relationship between order of acquisition and transmission of TYLCV and ToMoV, these results support the hypothesis that vector-virus interactions during acquisition by and/or circulation through the vector influence transmission outcomes.

### How viruses are acquired influences infection of new host plants

Virus-virus interactions occur in specific plant tissues and at specific time points during the infection cycle, but these interactions have both spatial and temporal dimensions that can be altered by virus-plant, vector-plant and vector-virus-plant interactions^42,43,65^. In our experimental treatments, temporal differences in transmission of the two viruses would be influenced by the timing and order in which the viruses were acquired from the source plants, the latent period, and whitefly settling behavior. Spatial effects on transmission were more likely in the mixed-cohort treatment (TYLCV+ToMoV) because individual whiteflies could inoculate only one virus and it is highly unlikely that any individuals shared a specific feeding site. In this scenario, temporal differences are also expected due to variation in whitefly settling behaviors that affect the amount of time before feeding and transmission begins.

Our results show that the ways in which two viruses are acquired can influence both the probability that they are transmitted (Table 3) and the occurrence of systemic infection in new hosts following transmission (Tables 4). Acquisition of both ToMoV with TYLCV by whitefly cohorts, regardless of order, decreased the number of TYLCV systemic infections in all co-acquisition treatments. TYLCV produced an antagonistic effect on total ToMoV systemic infections only when both viruses were acquired from the same plant (Mixed) or when TYLCV was acquired first (TYLCV»ToMoV) (Table 4). In the Mixed treatment, vector cohorts acquired and transmitted ToMoV alone and together with TYLCV to artificial diet, but no single systemic infections of ToMoV VCSIPS were observed (Tables 2, 3 & 4). Although how the acquisition scenarios altered systemic plant-infection outcomes is not clear, a plant-mediated effect is likely because the probability of infection after co-inoculation (Table 4) did not always reflect the observed probability of transmission to artificial diet (Table 3). Differences we observed between virus detection in plants versus artificial diet may have been influenced by the detection limits of qPCR in sucrose sachets in which low virus copy numbers may be present. It is also possible that antagonistic interactions during early virus replication cycles negatively impacted replication in new hosts. Begomovirus infection induces host plant defenses, plant signaling pathways and diffusible signals, and alters cell cycles as well as other biological processes^54^. Our findings raise questions about the direction of diffusible signals across different cell types and between locations where initial infection occurs.

A vector-mediated induction of plant responses may also have influenced virus accumulation because co-acquisition and co-inoculation did not similarly affect the probabilities of systemic infection or virus titers across treatments. Vector and virus induced changes in affected plants may also produce additive, synergistic or antagonistic effects on one another. Transcriptional and proteomic differences are induced by virus infection and vector settling and feeding; a recent study of ToMoV showed that over time virus infection may reverse vector induced changes in the proteome of Lanai tomatoes^65^. Vector and virus induced changes can alter viral fitness and can cause chemical and physiological modifications that may influence vector behavior and fitness^86,87^.

Past studies measured titers in co-infected plants to examine competition and fitness of the replicating viruses, but we are unaware of any studies quantifying the virus titers in VCSIPs. These results suggest that co-inoculation of two viruses may impact virus fitness parameters related to transmission even when both viruses do not persist in the inoculated plant. Our results show that accumulation of the viruses in plants was influenced by the acquisition scenario treatments in both VCSIPs (Figure 1D) and co-infected plants (Figure 1C). Co-inoculation with TYLCV reduced virus accumulation of ToMoV in VCSIPs from TYLCV»ToMoV, ToMoV»TYLCV and TYLCV+ToMoV treatments when compared to plants inoculated with only one virus (Figure 1D). No statistical differences among TYLCV titers in VCSIPs were detected, but VCSIPs in all co-acquisition scenario treatments had numerically higher titers compared to single infection treatments (Figure 1D). Quantification of virus titers in VCSIPs was prompted by reports of highly variable outcomes of virus-virus interactions in the literature^3^. No unexpected differences in visual symptoms were observed and symptoms followed expectations based on infection status (ToMoV, TYLCV or co-infection) (Supplemental Figure 5).

## Conclusion

Overall, these results support the idea that virus persistence and accumulation in new hosts involves complex interplay between vectors, viruses and host plants during virus acquisition, circulation through the vectors, and inoculation into new host plant cells^88^. However, our findings do not provide an assessment of the relative impacts of virus-virus interactions that occur in the vector, virus-virus interactions that occur in the host plant, or vector-plant interactions during acquisition or transmission on propagation of co-acquired viruses. Virus-virus-plant interactions that influence fitness components associated with acquisition, transmission, and infection of new hosts have the potential to greatly alter the epidemiology of viruses by changing their abundance and distribution in natural and managed ecosystems. The finding that co-inoculation may influence virus fitness in VCSIPs is also intriguing given the diversity of vectors and viruses that have overlapping geographic and host ranges. Future studies are needed on the mechanisms occurring at the cellular, physiological, organismal, population and species levels to better understand the epidemiological and evolutionary significance of virus_1_-virus_2_-vector-plant interactions.

## Materials and Methods

### Whitefly Colony

In 2016, a whitefly colony was established from a population in Auburn, AL and reared in a greenhouse on *S. melongena* L., ‘Pinstripe Hybrid’ (Park Seed, Greenwood, SC), a dwarf eggplant variety that is non-host for ToMoV or TYLCV. Whiteflies were identified as *B. tabaci* MEAM1 by sequencing a partial mitochondrial COI gene^89^ (data not shown). New whitefly cohorts were initiated every 7-14 days to provide the four-to-seven-dayold adult whiteflies used in the experiment.

#### Inoculation of Plant with Infectious Clones

To standardize the virus populations of TYLCV, ToMoV, and co-infections, infections were initiated using clones corresponding to partial tandem dimers of ToMoV A (pNSB1906), ToMoV B (pNSB1877), TYLCV (pNSB1736), or pMON721 (empty vector) transformed into *R. radiobacter*^63^. All virus inoculations (rhizobacterium and whitefly) were conducted using two true-leaf *S. lycopersicum* L., variety ‘Florida Lanai’ planted in 11.5 cm^2^ pots^64^. Twenty plants were inoculated for each infection status: TYCLV, ToMoV and co-infections. Five plants were mock inoculated with the empty vector as a negative control. Rhizobacterium-inoculated plants were used for vector acquisition at 28 days after inoculation (dpi) because previous reports showed that TYLCV and ToMoV titers are highest at 10-24 dpi and decline by 31 dpi in ‘Lanai’ tomatoes^64,90^. All experiments were conducted in a growth chamber (Percival Scientific Inc.) equipped with supplemental lighting from two strips of plant Grow LED lights (Litever®) and set at 25±3ºC and a 16:8 h light/dark cycle.

### Vector Acquisition and Co-inoculation of Viruses

Treatments compared different acquisition and inoculation scenarios that mimic the ways in which whiteflies can acquire and transmit single or co-infections of TYLCV and ToMoV (Table 1). The durations of AAPs and IAPs were chosen based on times reported to maximize transmission rates of these viruses^82,83,91^. Each treatment was replicated six times with each replicate using a different inoculated source plant. Each replicate of treatments 1-3 used one singly or doubly infected source plant, and each replicate of treatments 4-6 used two singly infected plants during the AAPs (Table 1). Infected plants were placed individually into a 32.5 cm × 32.5 cm × 32.5 cm cage (Product: BugDorm4S3030D MegaView Science Co.) with 500 non-viruliferous whiteflies for a total 48h AAP for each treatment. Three replications of each treatment were initiated the same day, and experiments were repeated twice for a total of six replications of each treatment. A non-infected mock plant served as a negative control throughout transmission experiments to confirm the whitefly colony was virus-free, monitor for unintended virus spread in incubators during experiments, and to use as negative controls for PCR and qPCR.

Following the AAP, cohorts of six female whiteflies were given a 48-h IAP on artificial diet or healthy tomato plants. To measure transmission to artificial diet, whitefly cohorts were aspirated into a 1.7-mL tube with a sucrose sachet made of parafilm (PARAFILM® M) stretched and folded thinly over the lid that contained 20 µL of a 20% sucrose solution. The lids were colored with a green Sharpie® (Sanford, L.P.) to artificially simulate the color of plant leaves and encourage whitefly feeding. After 48 h, the 1.7-mL tubes containing the whitefly cohorts and sucrose sachets were stored at -80□C (Revco UxF freezer, Thermo Fisher Scientific Inc) until further analysis. Due to low numbers of whiteflies during the first replications, sucrose sachet samples were only available from the second experimental replicate. Five sucrose sachets were inoculated in each of the three replicates per treatment (15 sucrose sachets per treatment in total). To measure transmission to tomato plants post-AAP, whitefly cohorts were aspirated into clip cages to confine them on the largest leaf of a healthy two-true-leaf tomato seedling. After the 48-h IAP, adult whiteflies were collected from the leaves and killed by freezing. At 14 dpi, a soil drench of imidacloprid (Admire® Pro, Bayer CropScience) was applied to kill whitefly nymphs. Plants were sampled at 28 dpi and virus infection was confirmed by PCR (described below). Ten plants were inoculated in each of the six replicates per treatment.

Overall lower survival of whiteflies on the co-infected plants reduced the sample sizes in the Mixed treatment. We note, however, that whitefly survival on rhizobacterium-inoculated plants coinfected with TYLCV and ToMoV was lower than on single virus-infected plants and the coinfected plants had an unusually strong plant odor (personal observation). To accommodate this, we infested the virus source plants in the Mixed treatment with a greater number of whiteflies to avoid confounding effects of vector mortality on observed transmission rates.

### Plant Tissue Sampling and DNA Extraction of Plants and Whiteflies

Leaf samples were collected from rhizobacterium-inoculated and whitefly-inoculated tomatoes at 28 dpi. A sterilized one-Hole Punch (6 mm in diameter) (Staples®) was used to take a composite sample comprised of two leaf discs from the terminal leaflet of the first fully expanded leaf below the apex of the plant, one leaf disc from terminal leaflet of the second fully expanded leaf, and one leaf disc from terminal leaflet of third fully expanded leaf. Samples from each plant were placed together in a 1.5-mL microcentrifuge tube with 3 mm glass beads (Millipore Sigma Darmstadt), flash frozen in liquid nitrogen, and stored at -80□. Before DNA extraction, frozen plant tissue samples were transported in liquid nitrogen to and from a Mini-BeadBeater™ (BioSpec Products, Inc.), where the samples were ground into a fine powder. DNA extractions were performed using the DNeasy® Plant Mini Kit (QIAGEN) following manufacturer’s instructions with minor revisions for eluting DNA using Buffer AE in a final volume of 60 µL.

Total DNA was extracted from cohorts of six whiteflies that fed on sucrose sachets using Qiagen® DNeasy Blood and Tissue Kit (QIAGEN) with the following modifications. Whiteflies were placed into 1.5-mL microcentrifuge tubes and 60 µL of Buffer ATL and 7 µL Proteinase K. Whiteflies were homologized into the liquid using a sterile 10-µL pipet tip to grind them against the sides of the tube. Volumes of the solutions were adjusted for low-tissue extractions using following volumes in the order in which they appear in the protocol: 100 µL Buffer AL, 100 µL 100% ethanol, 250 µL Buffer AW1, 250 µL Buffer AW2, and 30 µL Buffer AE repeated twice to yield a total volume of 60 µL.

### Virus Detection in Plant Samples Using PCR

Virus detection in plant samples was conducted using PCR according to the methods of Rajabu^64^ using primer pairs TYLCV15for/TYLCV15-rev (ca. 257 bp) and ToMoV pNSB1/ToMoV pNSB2 for ToMoV-A (ca. 239). Methods of Gong^90^ were used to amplify ToMoV-B-Fw/ToMoV-B-Rv (ca. 205). Amplified products and a 100-Kb DNA ladder (New England Biolabs Inc.) were visualized using agarose gel electrophoresis.

### Quantitative PCR (qPCR) of Viral DNA in Plant, Whitefly and Artificial Diet

Titers of TYLCV, ToMoV-A and ToMoV-B were quantified from virus source plants, whitefly cohorts, sucrose sachets, and whitefly-inoculated plants. Aforementioned primers were used for quantification of TYLCV and ToMoV-A, while ToMoV-B was quantified using the primer pair, ToMoV-B-Fw and ToMoV-B-Rv^90^. Total plant DNA and whitefly DNA were measured using a Nanodrop 2000 Spectrophotometer (Thermo Fisher Scientific Inc.) and a Qubit 3.0 Fluorometer (Thermo Fisher Scientific Inc), respectively, following manufacturers’ protocols for dsDNA and ssDNA HS (High Sensitivity) DNA Quantification. Plant DNA was standardized to 25 ng/5 µL of total DNA per qPCR reaction, and total DNA of whitefly cohorts was standardized to 1 ng/5 µL. Sucrose solution from sachets were added without modification in 5-µL volumes to each reaction. The amplification reactions had a total volume of 20 µL containing 5 µL of template DNA, 10 µL iTaq™ Universal SYBR® Green Supermix (Bio-Rad Laboratories, Inc.), 1 µL of 10-µM virus forward primer, 1 µL 10-µM virus reverse primer, and 3 µL sterile H2O. The qPCR analyses were performed using C1000 Touch™ thermocycler (Bio-Rad Laboratories, Inc., Hercules, CA) under conditions described in Gong^90^. Absolute quantification of titers was determined for each sample by comparison to a standard curve as described in Rajabu^64^. All samples and standards were run in triplicate.

Viral titers were quantified from five whitefly cohorts and their respective sucrose sachets from each replicate. Co-inoculation of TYLCV and ToMoV by whiteflies in the co-infection scenario treatments resulted in both singly and co-infected plants. Titers were quantified in at least one infected plant from each replicate and infection status (TYLCV, ToMoV, co-infected).

## Statistical analysis

Statistical analyses were performed using SAS PROC GLIMMIX version 9.4 (SAS Institute, Cary, NC). Tukey’s tests were used to perform LS means comparisons using a significance value of *P* <0.05. There were no significant differences among experimental replicates, so data were pooled for analysis. The proportion of samples testing positive for single infections of TYLCV, single infections of ToMoV, co-infections, and the total of single and co-infections for each virus were compared among treatments in separate analyses using a binary distribution. Presence or absence of the viruses was determined from PCR data from plant samples and qPCR data from whitefly and sucrose sachets. Three separate analyses compared titers of TYLCV, ToMoV-A and ToMoV-B among treatments. A fourth analysis only included samples that contained both viruses. These analyses were conducted separately for whitefly cohorts, sucrose sachets and plants using the main effects of treatment and virus/viral DNA component, and their interaction term. Titer data were log_10_ transformed and analyzed using a Gaussian distribution. Analyses of titer data for whitefly inoculated plants were conducted in three separate analyses based on infection status: TYLCV, ToMoV, and coinfection.

## Acknowledgements

Funding for this research was provided by National Science Foundation grant number OISE 1545553 (PI: L.H.B, co-PI G.G.K., senior scientist A.L.J.), and a graduate research assistantship provided by the Department of Entomology and Plant Pathology at Auburn University. Hatch projects supporting this research include AL-1021180, NC06640, and NC02784. The authors would like to thank Adam Kesheimer, Dr. Fredericka Hamilton, Anna Dye, John Mahas, Dr. Benard Mware, Dr. Kassie Conner, Dr. John Beckmann, and Dr. Nannan Liu for their technical support and assistance during these experiments.

## Author Contributions

A.A.M. assisted with the conceptualization of experiments, conducted experiments, statistical analysis, and wrote an early draft of the manuscript. A.L.J. provided funding and resources, conceptualized experiments, supervised research, assisted with statistical analysis, and finalized the manuscript draft. G.G.K. assisted with statistical analysis and editing of this manuscript. L.H.B. assisted in editing of this manuscript. All authors reviewed the manuscript.

## Data Availability

Data are available upon request to the corresponding author.

## Research Involving Plants

The plant collection and use was in accordance with all the relevant guidelines.

