## Supplemental Figure for "Vector acquisition and co-inoculation of two plant viruses influences transmission, infection, and replication in new hosts"

Log 10 Virus Titer

| Virus or Viral Component | TYLCV | ToMoV | Co-Infected |
| --- | --- | --- | --- |
| TYLCV | 20.3897 |  | 20.3779 |
| ToMoV-A |  | 19.3336 | 19.4749 |
| ToMoV-B |  | 19.9587 | 20.223 |

**Supplemental Figure 1.** The average log_10_ titers of TYLCV, ToMoV-A and ToMoV-B components in rhizobacterium-inoculated plants 28 days after inoculation; plants were used in acquisition access periods (AAPs). LS means comparisons were performed using Tukey’s method at *P*=0.05 by modeling the interaction between virus or viral component (TYLCV, ToMoV-A, or ToMoV_B) and infection status (TYLCV singly, ToMoV singly or co-infected). There were significant differences among virus titers (F _5, 65_ = 7.63, *P*<0.0001). Bars with different letters indicate significantly different values.

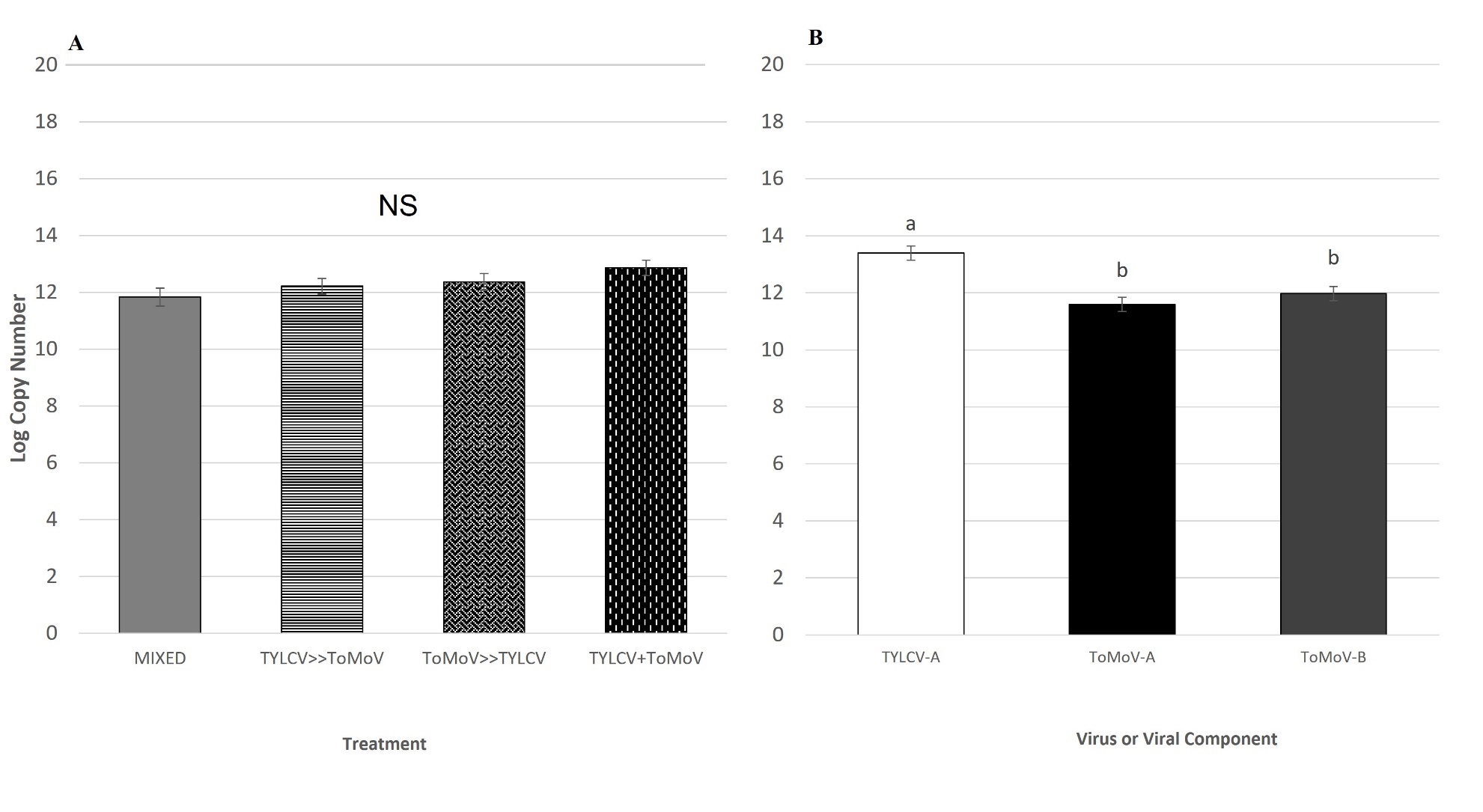

**Supplemental Figure 2.** The average log_10_ titers of TYLCV, ToMoV-A or/and ToMoV-B quantified in whiteflies. LS means comparisons on the main effects of treatment (A), virus titer (B), and their interaction (presented in main text, Figure 2A) were conducted using Tukey’s method at *P*=0.05. NS indicates no significant differences; bars with different letters indicate significantly different values.

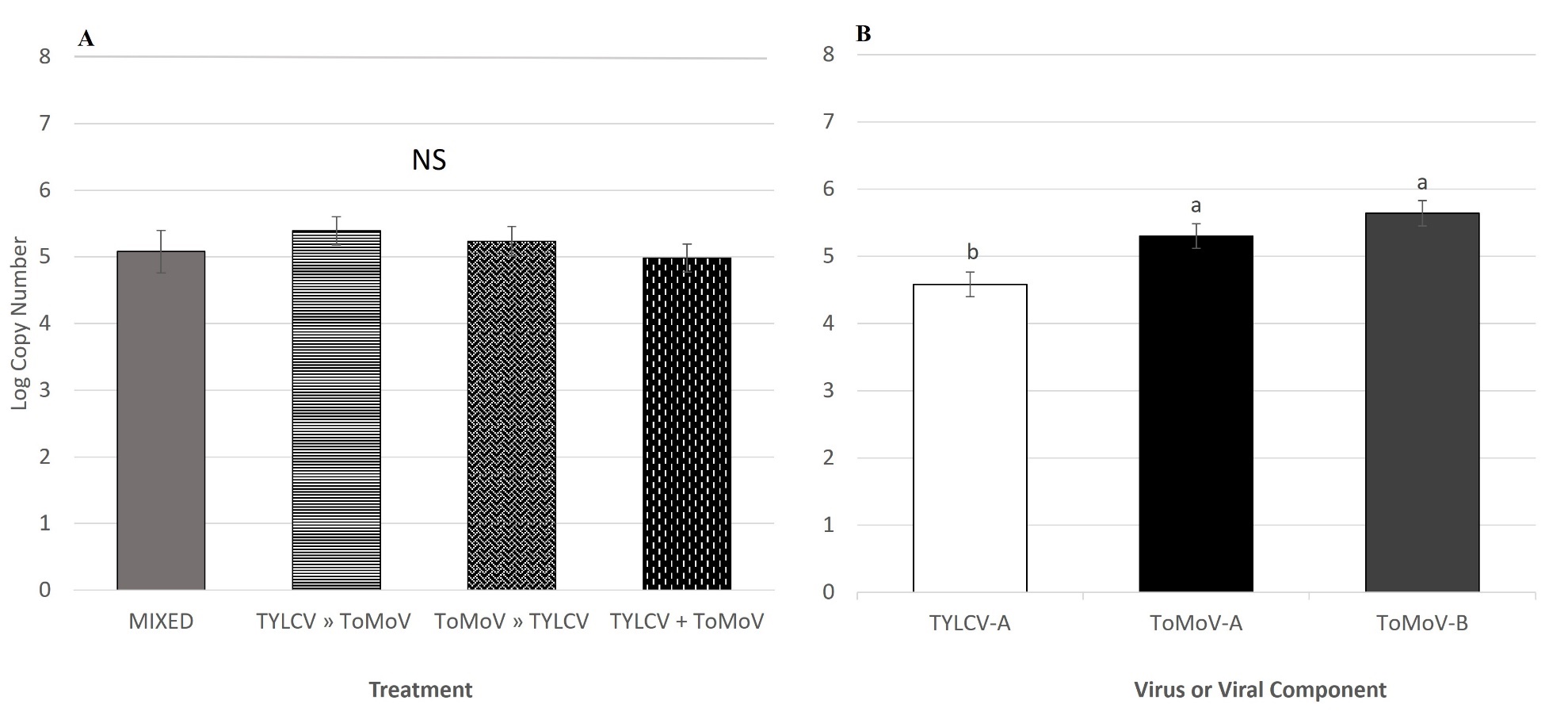

**Supplemental Figure 3.** The average log_10_ titers of TYLCV, ToMoV-A or/and ToMoV-B quantified in sucrose sachets. LS means comparisons on the main effects of (A) treatment, (B) DNA component and their interaction (presented in main text, Figure 2B) were conducted using Tukey’s method at *P*=0.05. NS indicates no significant differences; bars with different letters indicate significantly different values.

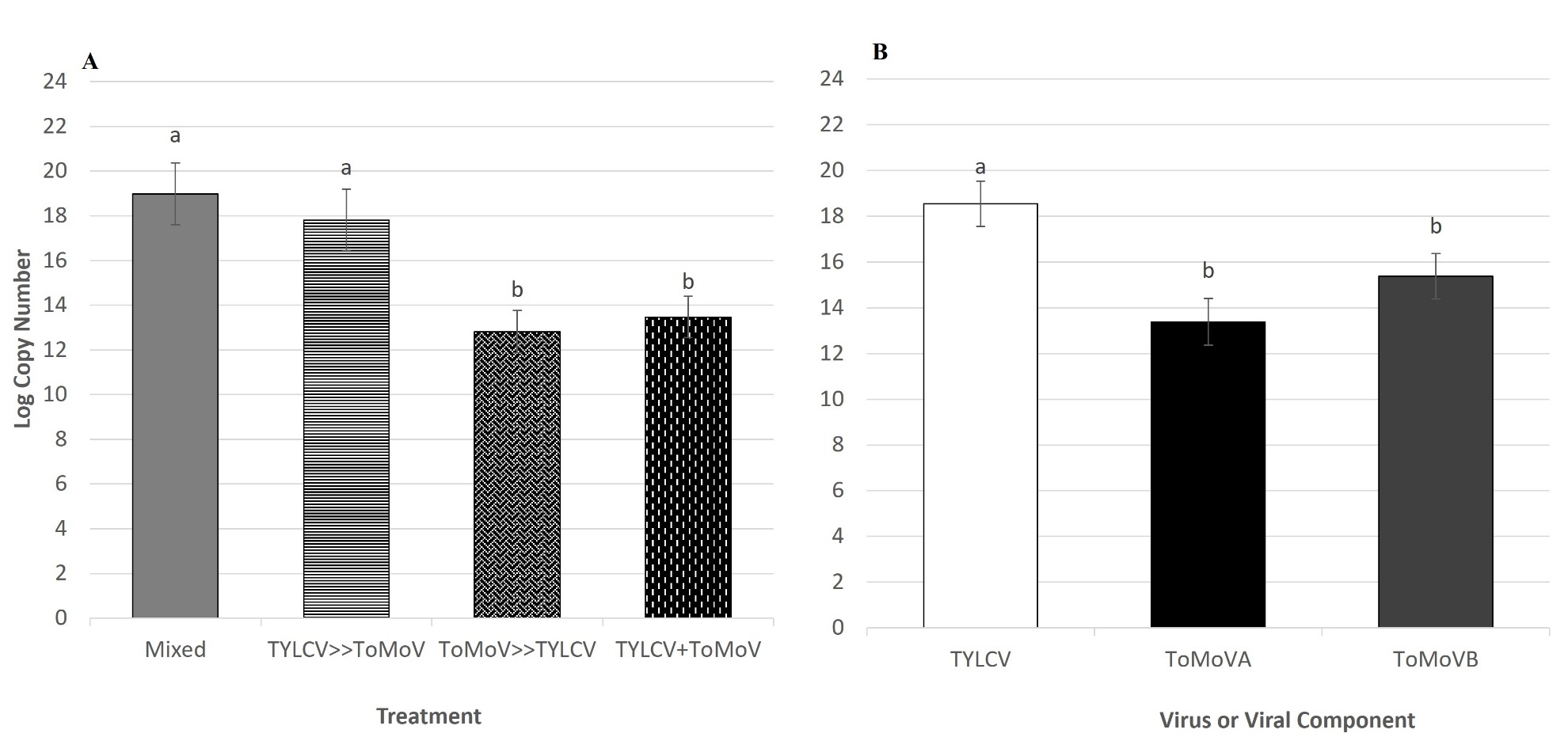

**Supplemental Figure 4.** The average log_10_ titers of TYLCV, ToMoV-A or/and ToMoV-B quantified in whitefly inoculated plants. LS means comparisons on the main effects of treatment (A), DNA component (B) and their interaction (presented in main text, Figure 2C) were conducted using Tukey’s method at *P*=0.05. Bars with different letters indicate significantly different values.

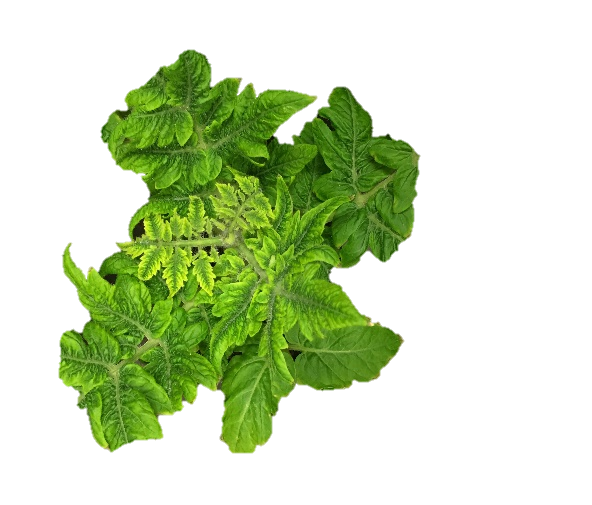

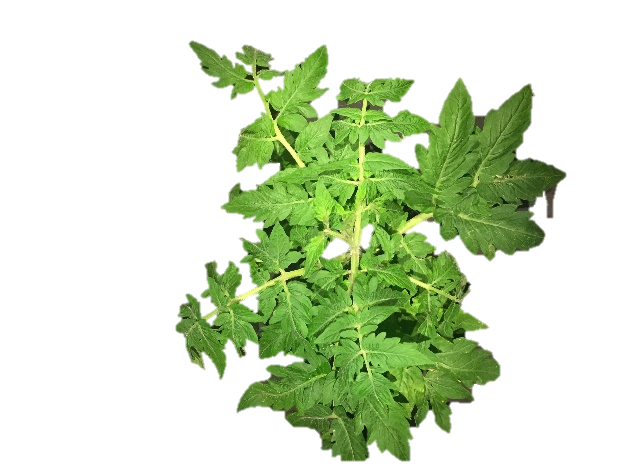

**A**

**B**

**C**

**D**

**Supplemental Figure 5.** Pictures of tomato plants 28 days post whitefly inoculation showing typical appearance of (A) non-infected control plants, and virus symptoms produced on (B) TYLCVinfected, (C) ToMoV-infected, and (D) co-infected (TYLCV & ToMoV) plants. Symptoms depicted were typical of all virus-infected plants in this study regardless of whether the plants were infected using rhizobacterium or whiteflies.
